# Rapidly cycling stem cells regenerate the intestine independent of *Lgr5^high^* cells

**DOI:** 10.1101/813402

**Authors:** Xiaole Sheng, Ziguang Lin, Cong Lv, Chunlei Shao, Xueyun Bi, Min Deng, Jiuzhi Xu, Christian F. Guerrero-Juarez, Mengzhen Li, Xi Wu, Ran Zhao, Xiaowei Liu, Qingyu Wang, Qing Nie, Wei Cui, Shan Gao, Hongquan Zhang, Zhihua Liu, Yingzi Cong, Maksim V. Plikus, Christopher J. Lengner, Bogi Andersen, Fazheng Ren, Zhengquan Yu

**Author notes:** These authors contribute equally (Xiaole Sheng, Ziguang Lin, Cong Lv). Co-corresponding authors: Address requests for reprints to: Zhengquan Yu, PhD, State Key Laboratories for Agrobiotechnology and Beijing Advanced Innovation Center for Food Nutrition and Human Health and, College of Biological Sciences, China Agricultural University, Yuanmingyuan West Rd. 2, Haidian District, Beijing, 100193, China,; Fazheng Ren, Beijing Advanced Innovation Center for Food Nutrition and Human Health, China Agricultural University, Beijing, China.

## Abstract

The +4 cells in intestinal crypts are DNA damage-resistant and contribute to regeneration. However, their exact identity and the mechanism underlying +4 cell-mediated regeneration remain unclear. Using lineage tracing, we show that cells marked by an *Msi1* reporter (*Msi1^+^*) are enriched at the +4 position in intestinal crypts and exhibit DNA damage resistance. Single-cell RNA sequencing reveals that the *Msi1^+^* cells are heterogeneous with the majority being intestinal stem cells (ISCs). The DNA damage-resistant subpopulation of *Msi1^+^* cells is characterized by low-to-negative *Lgr5* expression and is more rapidly cycling than *Lgr5^high^* radio-sensitive crypt base columnar stem cells (CBCs); they enable fast repopulation of the intestinal epithelium independent of CBCs that are largely depleted after irradiation. Furthermore, relative to CBCs, *Msi1^+^* cells preferentially produce Paneth cells during homeostasis and upon radiation repair. Together, we demonstrate that the DNA damage-resistant *Msi1^+^* cells are rapidly cycling ISCs that maintain and regenerate the intestinal epithelium.

## Introduction

The intestinal epithelium is a single-layer tissue organized into repetitive crypt-villus units. The cells that drive homeostatic intestinal renewal reside at the bottom of the crypt and move upwards toward the villus tip, where they eventually die – a process referred to as the conveyer-belt model (Heath, 1996). The intestinal epithelium undergoes rapid turnover, with the majority of epithelial cells replaced in three to five days in mice (Heath, 1996). The rapid turnover of intestinal epithelial cells renders them sensitive to irradiation. Consequently, patients undergoing radiation therapy to the abdomen, pelvis, or rectum develop acute enteritis, displaying symptoms that include pain, bloating, nausea, fecal urgency, diarrhea and rectal bleeding (Stacey & Green, 2014). Mucosal healing is critical for the remission of DNA damage-induced enteritis. Therefore, elucidating the cellular mechanisms of mucosal healing is necessary to develop new therapies for post-radiation enteritis.

Intestinal stem cells (ISCs), which reside within the proliferative compartment of crypts, are responsible for both intestinal homeostasis and epithelial regeneration after radiation exposure (Barker, 2014, Li & Clevers, 2010). Multiple studies have shown the existence of two functionally distinct ISC populations (Barker, 2014, Gehart & Clevers, 2015, Li & Clevers, 2010): mitotically active *Lgr5^high^* ISCs, commonly known as crypt base columnar stem cells, or CBCs (Barker, van Es et al., 2007, Sato, Vries et al., 2009), and a more dormant, reserve ISCs, defined as +4 cells due to their location within crypts (Li, Yousefi et al., 2014, Montgomery, Carlone et al., 2011, Powell, Wang et al., 2012, Sangiorgi & Capecchi, 2008, Takeda, Jain et al., 2011, Tian, Biehs et al., 2011). Although CBCs mainly function to maintain physiological homeostasis of intestinal epithelium (Barker et al., 2007, Sato et al., 2009), they are also thought to be indispensable for epithelial regeneration (Metcalfe, Kljavin et al., 2014). *In vitro,* a single *Lgr5^high^* CBC can form a mini-gut structure that contains all mature intestinal cell types (Sato et al., 2009). Therefore, CBCs have been proposed to be *bona fide* ISCs. In contrast, considerable controversy exists regarding the precise identity of +4 cells and their lineage relationship to CBCs. It remains unclear whether +4 cells are *bona fide* ISCs. Several markers of +4 cells, including *Bmi1*, *mTert*, *Hopx* and *Lrig1*, have been identified by *in vivo* lineage tracing, either by knockin of CreER into the gene or by randomly-integrated transgenesis (Barker, 2014, Montgomery et al., 2011, Sangiorgi & Capecchi, 2008, Takeda et al., 2011, Tian et al., 2011). In contrast to CBCs, +4 cells are resistant to DNA damage, and are activated to promote epithelial regeneration upon radiation-induced CBC depletion. Therefore, +4 cells are thought to be reserve ISCs, and their cell cycle quiescence has been proposed to be the main source of their radioresistance. The primary evidence for +4 cells’ quiescence is that the +4 position corresponds to label retaining cells (Potten, Kovacs et al., 1974, Potten, Owen et al., 2002) and that Bmi1-, Hopx- or Lrig1-marked +4 cells undergo slower kinetics of lineage tracing in comparison to *Lgr5*-expressing ISCs (Powell et al., 2012, Takeda et al., 2011, Yan, Chia et al., 2012). Further evidence is that *Hopx^CreER^* cells were shown to reside in G0 (Li, Nakauka-Ddamba et al., 2016). However, three independent studies have demonstrated that label-retaining cells are in fact terminally differentiated Paneth cells or secretory progenitors (Buczacki, Zecchini et al., 2013, Li et al., 2016, Steinhauser, Bailey et al., 2012). Additionally, it is worth mentioning that the primary DNA damage repair pathway in quiescent stem cells—non-homologous end joining (NHEJ)—is error-prone (Mohrin, Bourke et al., 2010), which is unfavorable for tissue repair, while homologous recombination (HR)-mediated acute DNA repair can only occur in cycling cells during late S and G2 phases (Moynahan & Jasin, 2010). Therefore, the identity of +4 cells and the mechanisms underlying +4 cell-mediated epithelial regeneration remain uncertain.

Here we generated an *Msi1^CreERT2^* allele for lineage tracing and observe that *Msi1^CreERT2^-*marked cells are enriched at the +4 crypt position, referred as *Msi1^+^* cells, and are resistant to DNA damage. Single-cell RNA sequencing (scRNA-seq) of *Msi1^+^* cells further revealed that a subset of S/G2-phase rapidly cycling stem cells, characterized by low-to-negative *Lgr5* expression, exhibit DNA-damage resistance and repopulate radiation-damaged epithelium independent of CBCs, which substantially differs from the classic theory that such +4 cells function as reserve stem cells, activate following irradiation to restore the depleted *Lgr5^high^* CBCs first, and then the nascent CBCs rapidly divide to repair damaged intestinal epithelium. Furthermore, we observed that *Msi1^+^* cells preferentially produce the Paneth lineage, relative to CBCs.

## Results

### An Msi1 reporter marks DNA damage-resistant intestinal +4 stem cells

Msi1 has been identified as a marker for ISCs, including both CBCs and +4 cells (Kayahara, Sawada et al., 2003, Li, Yousefi et al., 2015). We first validated Msi1 expression pattern in CBCs and +4 cells at the protein level (**Fig EV1A**). At the RNA level, *Msi1* expression was the strongest in +4 cells (**Fig EV1B**). To track the fate of *Msi1*-expressing cells within intestinal epithelium, we generated a tamoxifen-inducible Cre (CreERT2) knock-in targeted just before the stop codon of the endogenous *Msi1* locus (**Fig EV1C**). We then crossed *Msi1^CreERT2^* mice with *R26^Lox-Stop-Lox-LacZ^* (*R26R^LacZ^*) reporter mice. Fifteen hours after one pulse of tamoxifen, X-gal staining showed that 15.7% of intestinal crypts were labeled and that *Msi1* reporter-marked cells were mainly located at the +4 position of intestinal crypts (n=472 crypts from 3 mice, **Fig 1A and B**), which we further corroborated in *Msi1^CreERT2^;R26^mTmG^* mice (n=99 crypts from 3 mice, **Fig 1C**). Thus, *Msi1^CreERT2^*-marked cells are largely positionally distinct from *Lgr5^high^* CBCs (**Fig 1A and B**).

**Figure 1.**
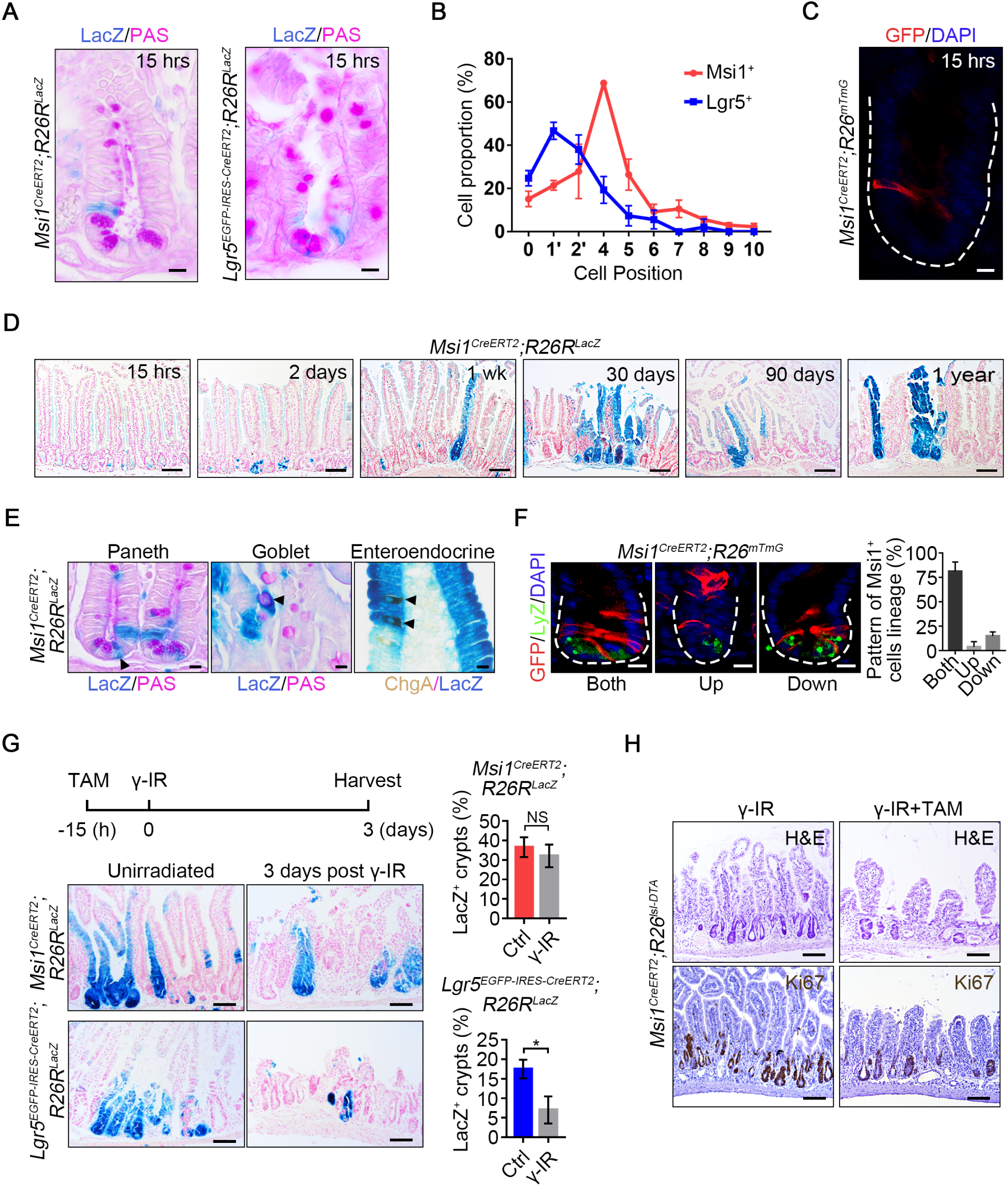
*Msi1* reporter-marked +4 ISCs in intestinal crypts. (A) Representative images of LacZ^+^ cells in *Msi1^CreERT2^;R26R^LacZ^*(302 crypts; n = 3 mice) and *Lgr5^EGFP-IRES-CreERT2^;R26R^LacZ^*(175 crypts; n = 3 mice) lineage-labeled small intestinal crypts fifteen hours after one single dose of 4 mg/25 g tamoxifen induction. Scale bar: 10 μm. (B) Quantification of the position of LacZ^+^ cells in intestinal crypts in Panel A. n = 3 mice at each time point. (C) Representative images of GFP^+^ cells in *Msi1^CreERT2^;R26^mTmG^* lineage-labeled small intestinal crypts fifteen hours after tamoxifen induction. Scale bar: 10 μm. (D) Low-magnification images of the LacZ^+^ ribbon in *Msi1^CreERT2^;R26R^LacZ^* lineage-labeled small intestine at different time points following tamoxifen induction. n ≥ 3 mice at each time point. Scale bar: 40 μm. (E) PAS staining and immunohistochemistry for ChgA in *Msi1^CreERT2^;R26R^LacZ^* lineage-labeled small intestine one week after tamoxifen induction. n = 3 mice. Scale bar: 10 μm. (F) Double immunofluorescence for GFP and lysozyme in *Msi1^CreERT2^;R26^mTmG^*(n=3 mice) lineage-labeled small intestinal crypts twenty-four hours after tamoxifen induction. The position of GFP+ cells is above the +4 position, referred to as “Up”; below the +4 position, referred to as “Down”; above and below the +4 position, referred to as “Both”. Quantification of the lineage pattern of *Msi1*-reporter positive cells. Scale bar: 10 μm. (G) Representative images of LacZ^+^ ribbons in *Msi1^CreERT2^;R26R^LacZ^* and *Lgr5^EGFP-IRES-CreERT2^;R26R^LacZ^* lineage-labeled small intestines four days after tamoxifen induction, or the mice were irradiated after fifteen hours of tamoxifen exposure, and harvested three days after 12 Gy γ-IR. n = 3 mice at each time point. Scale bar: 10 μm. Quantification of LacZ^+^ ribbons under the indicated conditions. (H) Depletion of *Msi1^+^* cells impaired intestinal epithelial regeneration. *Msi1^CreERT2^;R26^lsl-DTA^* mice irradiated fifteen hours after tamoxifen induction and then harvested three days after γ-IR. H&E and immunohistochemistry for Ki67 under the indicated conditions. Scale bar: 100 μm. In G, data represent the mean value ± SD. NS, not significant; \**P*<0.05 (Student’s t-test).

Two days after tamoxifen induction, most labeled crypts contained 3 to 7 cells exhibiting β-galactosidase activity (**Fig 1D**). One week after induction, X-gal staining became more widespread (**Fig 1D**), and the labeling cells included differentiated cell lineages – Paneth, goblet and enteroendocrine cells (EECs) (**Fig 1E**). The number of fully labeled crypt-villus ribbons increased over time (**Fig 1D, and EV1D and E**), and *Msi1* reporter marked progeny existed for at least 1 year (**Fig 1D**). Next, we sought to examine how *Msi1* reporter-marked cells give rise to distinct cell lineages. We quantified the positions of labeled cells one day after tamoxifen induction, a timepoint when newly generated cells are emerging, and found that the majority of labeled cells move both upwards and downwards relative to +4 position (**Fig 1F**). This distribution suggests that *Msi1* reporter-marked cells concomitantly give rise to distinct lineages, including CBCs, Paneth cells and villus cells. The expression of CBC markers and Wnt target genes (*Lgr5*, *Ascl2, Axin2, Sox9,* and *Olfm4*) in *Msi1^CreERT2^*-marked cells was similar to that of cells marked with *Hopx^CreERT2^*, a well-established marker of +4 ISCs (Takeda et al., 2011), and distinct from that of *Lgr5^high^* CBCs (**Fig EV1F and G**). Collectively, these data demonstrate that *Msi1* reporter-marked cells are primarily located above the crypt base and the CBC compartment and exhibit multipotent stem cell properties.

To examine the DNA damage response by *Msi1^+^* cells, we exposed *Msi1^CreERT2^;R26R^LacZ^ and Lgr5^EGFP-IRES-CreERT2^;R26R^LacZ^* mice to 12 Gy of ionizing radiation (γ-IR), fifteen hours after a single pulse of tamoxifen. After radiation exposure, the number of LacZ^+^ ribbons produced by *Msi1^+^* cells was similar to what we observed during homeostasis, while the number of LacZ^+^ ribbons from *Lgr5* reporter-marked cells was strongly reduced (**Fig 1G**). Lineage-tracing analysis in *Msi1^CreERT2^;R26^mTmG^* mice also demonstrated a robust repopulating capacity of *Msi1^+^* cells after exposure to γ-IR (**Fig EV1H**). The findings indicate that *Msi1^+^* cells are radio-resistant, able to survive γ-IR and repopulate the damaged epithelium. Moreover, depletion of *Msi1^+^* cells significantly impaired intestinal epithelial regeneration following γ-IR. (**Fig 1H and EV1I**). Taken together, these findings suggest that *Msi1^+^* cells are DNA damage-resistant ISCs with the capacity to repopulate γ-IR-damaged epithelium.

## *Msi1^+^* cells are a heterogeneous population

Next, we sought to better characterize the identity of *Msi1^+^* cells using single-cell RNA-sequencing (scRNA-seq) analysis. GFP-labeled cells from *Msi1^CreERT2^;R26^mTmG^* mice were sorted fifteen hours after tamoxifen induction, and subjected to scRNA-seq (**Fig EV2A**). Unsupervised clustering (Duo, Robinson et al., 2018) identified nine distinct cell clusters (**Fig 2A**). We utilized the differentially expressed gene signatures to assign putative cell type identities to these clusters (**Fig 2B-D, and EV2B and C**). Cluster H15h-C1 (at homeostasis, traced for 15 hours) is enriched in cells expressing the highest levels of ISC marker gene *Lgr5*, as well as several other ISC marker genes, namely, *Gkn3, Ascl2, Olfm4, Jun, Pdgfa* and *2210407c18Rik* (**Fig 2C and EV2C**). Thus, H15h-C1 cells were defined as *Lgr5^high^* ISCs. Clusters of H15h-C2 and H15h-C3 cells have low or negative *Lgr5* status, but concomitantly express the ISC marker genes *Igfbp4, Ascl2* and *Hopx* (**Fig 2C**), on which basis they are classified as *Lgr5^low/neg^* ISCs. In comparison to H15h-C2 cells, H15h-C3 cells highly express G2/M-phase marker genes (**Fig 2D**). Consistently, single-cell consensus clustering analysis (Kiselev, Kirschner et al., 2017) of clusters H15h-C1, -C2 and -C3 revealed higher similarity between H15h-C2 and H15h-C3 cells, relative to H15h-C1 ISCs; and further divided H15h-C1 cells into two sub-clusters (**Fig 2E**). Cluster H15h-C4 cells are also enriched for G2/M phase marker genes (**Fig 2D**), and principal component analysis (PCA) analysis shows that these cells are intermediate between ISCs and enterocytes (ECs) (**Fig 2F**). Thus, H15h-C4 cells were identified as EC precursor cells (EPs). Surprisingly, the smaller clusters H15h-C5 through H15h-C9 were characterized as differentiated cells – ECs, goblet cells, Paneth cells, EECs and tuft cells (**Fig EV2B**). These differentiated cells are likely the early-differentiated progeny of initially labeled +4 cells produced over the 15-hour period, suggesting that +4 cells have started the differentiation program.

**Figure 2.**
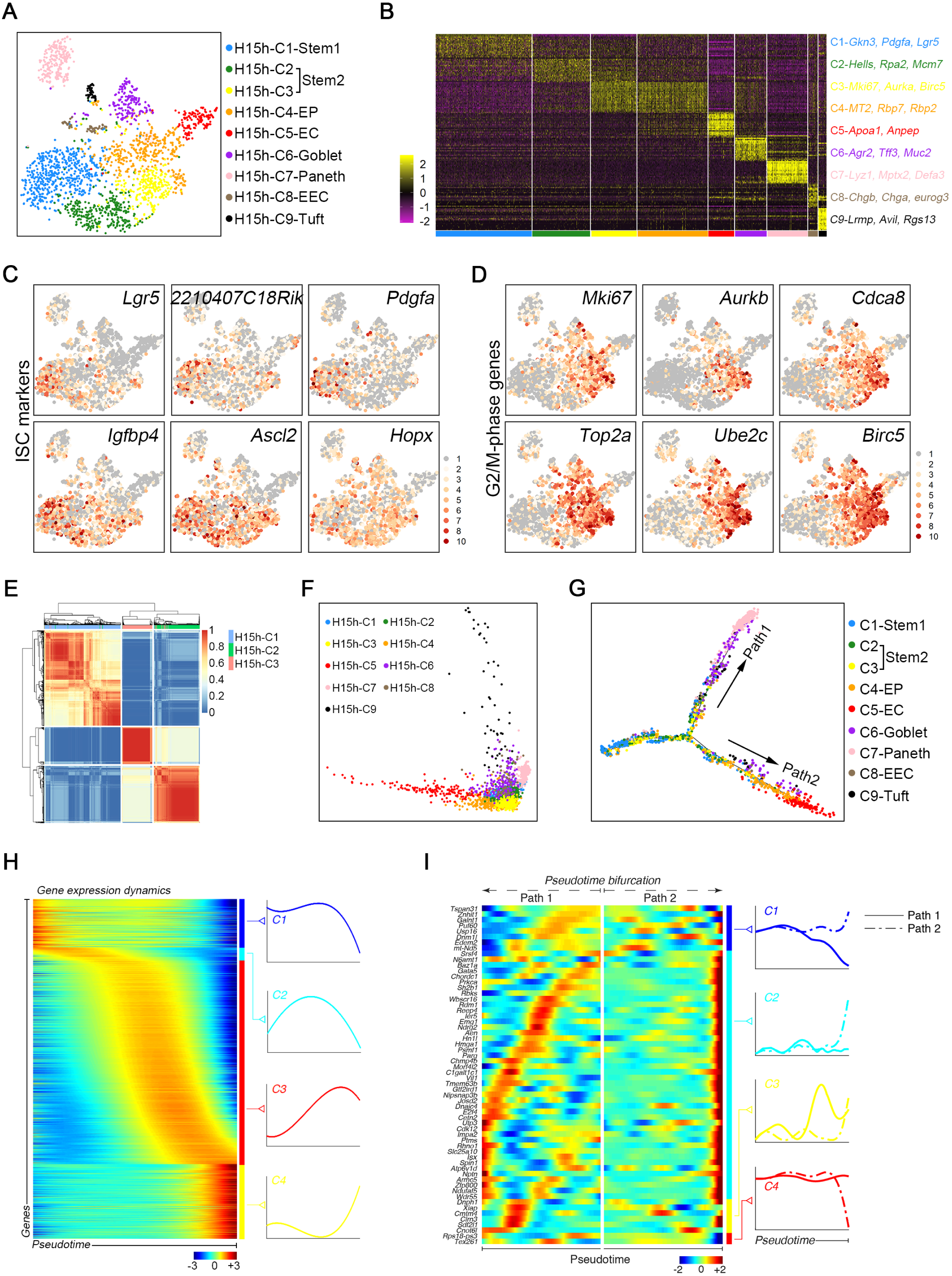
*Msi1^+^* cells are a heterogeneous population. (A) A t-SNE plot revealed cellular heterogeneity, with nine distinct clusters of *Msi1^+^* cells sorted from *Msi1^CreERT2^;R26^mTmG^* mice fifteen hours after tamoxifen induction. The general identity of each cell cluster is defined on the right. (B) Heatmap of differentially expressed genes in each cluster. Selected signature genes are shown on the right. (C and D) Feature plots of expression distribution for ISC (C) and G2/M phase (D) marker genes. Expression levels for each cell are color-coded. (E) Single-cell consensus clustering (SC3) analysis showing the correlation of H15h-C1 to H15h-C3. (F) PCA showing the association of distinct cell clusters. (G) Pseudotime ordering on *Msi1^+^* cells arranged them into a major trajectory with two branches of absorptive and secretary differentiated cells. (H) scEpath analysis performed on pseudotime along the trajectory from stem cells to differentiated cells, identifying four gene clusters (C1-C4) of pseudotime-dependent genes. (I) scEpath analysis identifying four gene clusters (C1-C4) of branching genes.

To understand the hierarchy among distinct cell clusters, we performed pseudo-temporal ordering of scRNA-seq data (Qiu, Hill et al., 2017). This analysis arranged most ISCs from the H15h-C1, -C2 and -C3 clusters into a major pseudotime trajectory that bifurcates toward ECs and differentiated secretory cells (**Fig 2G and EV2D**). Consistent with cluster identity attribution, ECs are preceded by EPs (H15h-C4 cells) in Path2 of the pseudotime (**Fig 2G**). A large number of genes were differentially expressed in cells along the pseudotime trajectory (**Fig 2H**). Among them, a number of “branching” genes were identified, which are potentially important for EC vs. secretory cell differentiation (**Fig 2I**). Taken together, these data indicate that the *Msi1^CreERT2^* allele marks a heterogenous population of cells, consisting primarily of ISCs and a small number of differentiated cells and residing along the two major differentiation trajectories.

### Rapidly cycling ISCs initiate epithelial regeneration

Next, we sought to define the initial cells that repopulate the epithelium after γ-IR exposure. We performed scRNA-seq on the progeny of *Msi1^+^* cells from *Msi1^CreERT2^;R26^mTmG^* mice two days after irradiation, a time point marking the initiation of epithelial regeneration (Kim, Yang et al., 2017). A minimal number (1-2) of proliferating cells exist in each crypt at this time point, followed by rapid proliferative expansion between 72 to 96 hours (**Fig EV3A and B**). Ten distinct cell clusters were identified, including clusters of stem cells, transition cells, EP-like cells, ECs, goblet cells, EECs, tuft cells and 3 distinct clusters of Paneth cells (**Fig 3A and B, and EV3C and D**). Importantly, the distribution of known ISC marker genes changed dramatically (**Fig 3C**). Compared to the distribution of *Msi1^+^* cells during homeostasis, the *Lgr5^high^* cell cluster was depleted two days after irradiation (**Fig 3C**). Consistently, the number of *Lgr5* reporter-marked cells becomes markedly reduced two days after irradiation or treatment with the DNA replication inhibitor and chemotherapeutic agent 5-fluorouracil (5-FU) (De Angelis, Svendsrud et al., 2006). In contrast, the number of *Msi1^+^* cells showed an increasing trend, albeit not significant, upon these treatments (**Fig 3D and EV3E**). Cluster IR2-C1 (2 days after irradiation) cells are identified as ISCs, as they strongly expressed ISC marker genes *Igfbp4* and *Ascl2* (**Fig 3C**). IR2-C2 cells were identified as a transition cluster due to their intermediate position between ISCs and differentiated cells (**Fig 3E**). IR2-C1 and IR2-C2 cells are enriched for genes functioning on DNA damage response (DDR) and cell survival (**Fig 3F and EV3F**), suggesting a strong DDR. In the Pseudotime trajectory, IR2-C1 and IR2-C2 cells are enriched at the starting point of the major branch, while IR2-C3 and IR2-C4 cells are enriched at the end of EC branch, with the remaining cells enriched at the end of the secretory/differentiated branch (**Fig 3G**). Consequently, few cells localize around the pseudotime bifurcation as compared with normal physiological conditions (**Fig 3G**). Importantly, IR2-C1 and IR2-C2 cells are rapidly cycling, while the other cells primarily reside in G0/G1 phase (**Fig 3H and EV3G**). These data suggest that IR2-C1 and IR2-C2 cells are rapidly-cycling ISCs that initiate epithelial regeneration.

**Figure 3.**
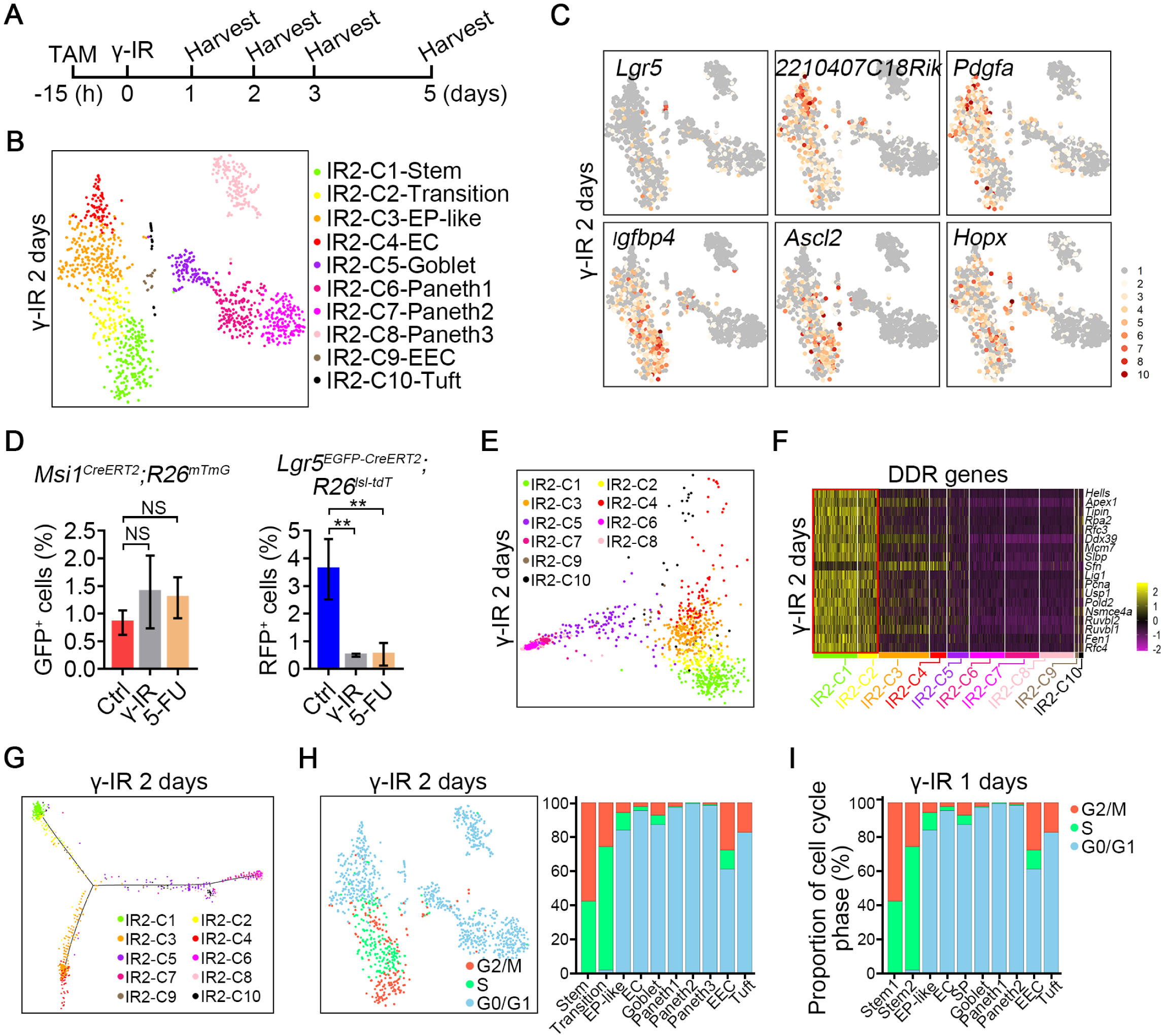
Rapidly cycling ISCs initiate intestinal epithelial regeneration. (A) Strategy of sample collection for scRNA-sequencing after irradiation. (B) A t-SNE plot revealed cellular heterogeneity with ten distinct clusters of *Msi1^+^* cell progeny from *Msi1^CreERT2^;R26^mTmG^* mice two days after γ-IR. The mice were pretreated with tamoxifen fifteen hours before irradiation. The general identity of each cell cluster is defined on the right. (C) Feature plots of expression distribution for ISC marker genes in t-SNE plots two days after irradiation. Expression levels for each cell are color-coded. (D) Quantification of *Msi1^+^* (n=3 mice) and *Lgr5^+^* (n=3 mice) populations two days after treatment with γ-IR or 5-FU. Mice were treated with γ-IR or two consecutive doses of 5-FU, and then induced by tamoxifen fifteen hours before sacrifice, as shown in Fig EV3E. \*\**P* < 0.01 (Student’s t-test). (E) PCA showing the association of distinct cell clusters two days after irradiation. (F) Heatmap of DDR genes in distinct clusters two days after γ-IR. (G) Pseudotime ordering of *Msi1^+^* cell progeny arranged distinct cluster cells into a major trajectory with two branches of absorptive and secretary differentiated cells two days after irradiation. (H) Cell cycle metrics of *Msi1^+^* cell progeny two days after γ-IR. t-SNE plot of the assigned cell cycle stages of *Msi1^+^* cell progeny. Cells in S phase are colored green, those in G2/M phase are red, and those in G0/G1 phase are blue. Proportions of cell cycle stages per cluster. (I) Proportions of cell cycle stages in each cluster one day after γ-IR. Cells in S phase are colored green, those in G2/M phase are red, and those in G0/G1 phase are blue. In D, data represent the mean value ± SD. NS, not significant; \**P*<0.05 (Student’s t-test).

Previous reports indicated that radio-resistant +4 cells are dormant (Montgomery et al., 2011, Powell et al., 2012, Yan et al., 2012). Therefore, on scRNA-seq we expected to see a certain number of quiescent ISCs (in G0/G1 phase), along with proliferatively active ISCs. In contrast, we found that approximately 42% of IR2-C1 cells were in S phase, and 58% were in G2/M phase. There were no G0/G1 phase IR2-C1 cells (**Fig 3H**). IR2-C1 cluster also strongly expressed proliferating cell marker genes (**Fig EV3G**). This data suggests that quiescent ISCs are lacking at this stage. Next, we considered that two days after irradiation might be too late to detect surviving quiescent ISCs. Thus, we performed scRNA-seq on the progeny of *Msi1^+^* cells from *Msi1^CreERT2^;R26^mTmG^* mice one day after irradiation, a time point when the majority of intestinal cells are undergoing cell death. Two clusters of ISCs were identified (**Fig EV3H**), and surprisingly, they also exhibited a highly proliferative state, with no cells in G0/G1 phase (**Fig 3I and EV3I**). Together, these data demonstrate that rapidly cycling, rather than quiescent ISCs initiate epithelial regeneration. It also raises the possibility that a population of rapidly cycling ISCs is resistant to and can survive irradiation exposure.

### Rapidly cycling Msi1^+^ ISCs survive from exposure to high dose of irradiation

To test whether rapidly cycling ISCs survive from exposure to high dose of irradiation, we labeled S-phase *Msi1^+^* cells using a 90 min pulse of EdU at 0.017mg/25 g body weight, which is insufficient to label all S phase cells (**Fig EV4A**), and then irradiated the mice. Indeed, we found that the labeled S-phase *Msi1^+^* cells survived γ-IR and divided, and the EdU signals diluted over time (**Fig 4A**). We then went back to the homeostatic condition and analyzed cell cycle phases of cells in intestinal crypts by quantifying the positions of PCNA^+^, EdU^+^ and pH3^+^ cells. Most cells in the +1 and +2 positions, which are usually considered to be *Lgr5^high^* CBCs (Barker et al., 2007), were in G1 phase (**Fig 4B**), while the cells from positions 4 to 6, referred to as *Lgr5^low/neg^*+4 cells with DNA damage resistance (Powell et al., 2012, Takeda et al., 2011), were in S or G2/M phases (**Fig 4B**). We then analyzed the division kinetics using dual BrdU/EdU labeling, and revealed that the average length of the cell cycle for +4 cells is 13.28 hours, while that of CBCs is 18.12 hours (**Fig 4C**). Furthermore, EdU labeling assay revealed that more *Msi1^+^* cells were in S-phase compared to *Lgr5^high^* CBCs (**Fig 4D and E**). Similar findings were also observed in *Hopx* reporter marked cells (**Fig 4D and E**). In agreement with those results, the majority of *Lgr5^low/neg^* ISCs from H15h-C2 and H15h-C3 scRNA-seq clusters reside in S and G2/M phases during homeostasis, while the majority of *Lgr5^high^* ISCs reside in G1 phase (**Fig 4F**). It has been shown that the signaling pathways regulating the DDR also activate during normal S-phase for genome integrity maintenance (Ben-Yehoyada, Gautier et al., 2007) and this property can increase cellular resistance to DNA damage. Accordingly, DDR genes are specifically enriched in rapidly-cycling H15h-C2 cells (**Fig 4G and EV4B**). H15h-C2 cells are enriched for genes functioning in cell survival and stress, which might facilitate cell survival after exposure to irradiation (**Figure EV4C**). The homologous recombination (HR)-mediated DNA repair only occurs in cycling cells at S- and G2-phases, enabling an accurate repair of DNA damage (Moynahan & Jasin, 2010). We found that the key components of HR-type repair such as *Rad51*, *Rad51ap1*, *Brca1*, *Brca2* and *Smc6* are highly expressed in the ISCs populations one and two days after irradiation **(Fig 4H and EV4D**), suggesting a strong HR-type repair response in S- and G2-phase ISCs. Taken together, these findings strongly suggest that the DNA damage-resistant *Msi1^+^* cells are more rapidly cycling than *Lgr5^high^* CBCs.

**Figure 4.**
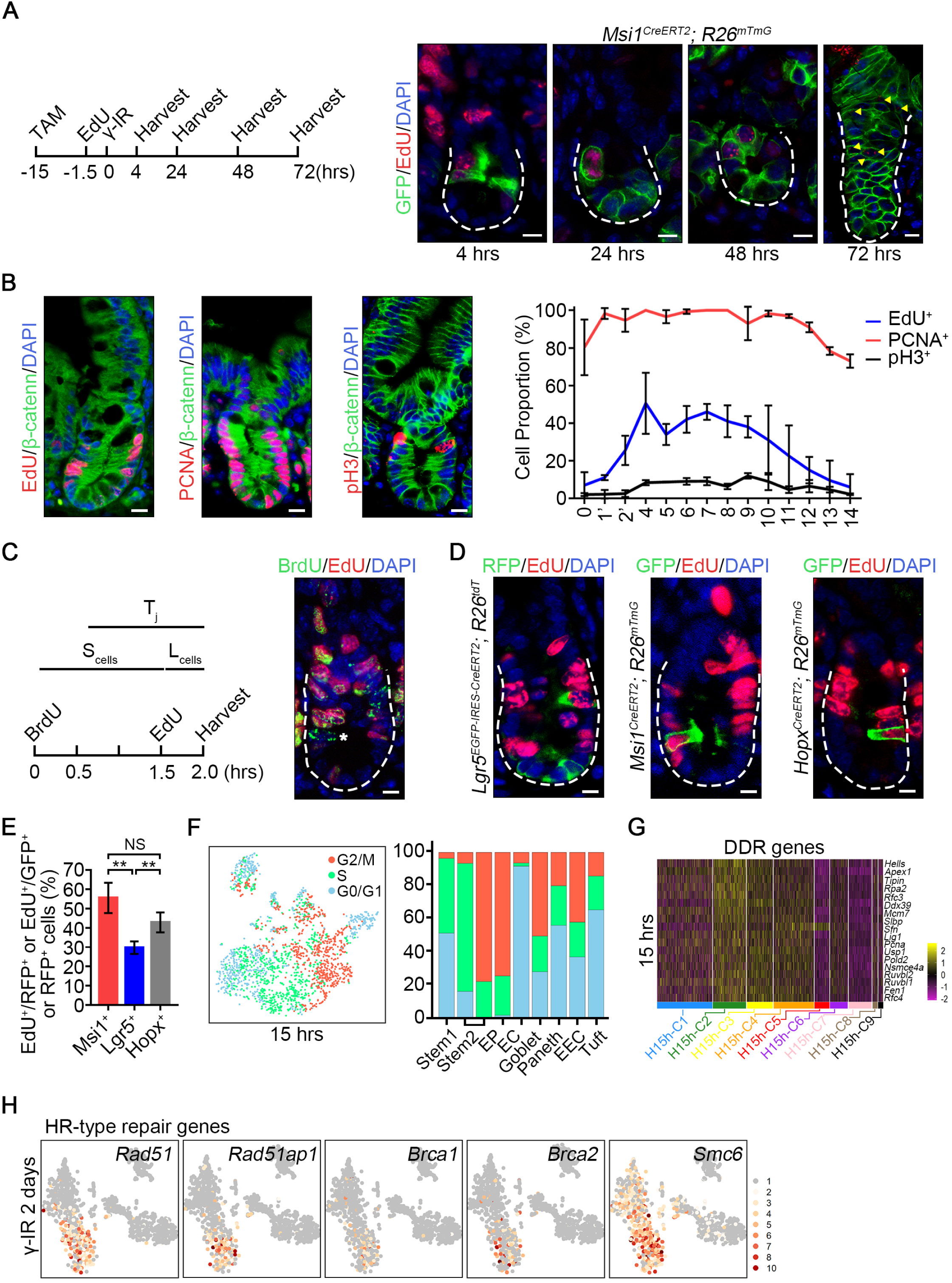
*Msi1^+^* cells are more rapidly cycling compared to CBCs. (A) Strategy of testing whether EdU labeled S-phase *Msi1^+^* cells survive from irradiation exposure. *Msi1^CreERT2^;R26^mTmG^* mice were treated by tamoxifen (TAM), and labeled using a 90 min pulse of EdU at 0.017mg/25 g body weight 13.5 hours after tamoxifen induction, and then irradiated fifteen hours after tamoxifen induction. Immunofluorescence for EdU and GFP in intestinal crypts at the indicated time points. Scale bar: 10 μm. (B) Immunofluorescence for EdU/β-catenin, PCNA/β-catenin, and phosphor-histone 3/β-catenin in the intestinal crypts of WT mice. Scale bar: 10 μm. Quantification of EdU^+^ (n = 100 Crypts), PCNA^+^ (n = 59 crypts) and pH3^+^ (n = 202 crypts) cells at the indicated position of intestinal crypts. (C) Schematics of the EdU and BrdU temporary space pulse method to calculate the average length of cell cycle times (left). Cells still in S phase during the labeled time were EdU^+^BrdU^+^, whereas cells that exited S phase were EdU^-^BrdU^+^, indicated by asterisks (right). (D) Immunofluorescence for RFP and EdU in intestines from *Lgr5^EGFP-CreERT2^;R26^lsl-tdT^* mice fifteen hours after tamoxifen induction, and immunofluorescence for GFP and EdU in intestines from *Msi1^CreERT2^;R26^mTmG^* and *Hopx^CreERT2^;R26^mTmG^* mice fifteen hours after tamoxifen induction. Scale bar: 10 μm. (E) Quantification of RFP^+^/EdU^+^ cells in *Lgr5^EGFP-CreERT2^;R26^lsl-tdT^* intestinal crypts (n = 268 cells) and GFP^+^/EdU^+^ cells in *Msi1^CreERT2^;R26^mTmG^* (n = 162 cells) and *Hopx^CreERT2^;R26^mTmG^* (n = 200 cells) intestinal crypts in Panel E. \*\**P*<0.01 (Student’s t-test). (F) Cell cycle metrics of *Msi1^+^* cells at homeostasis. t-SNE plot of assigned cycling stages on *Msi1^+^* cells. Cells in S phase are colored green, those in G2/M phase are red, and those in G0/G1 phase are blue. Proportions of cell cycle stages per cluster. (G) Heatmap of DDR genes in distinct clusters of *Msi1^+^* cells. (H) Feature plots of expression distribution for the key genes functioning in HR-type DNA damage repair two days after irradiation. Expression levels for each cell are color-coded. In E, data represent the mean value ± SD. \*\**P*<0.01 (Student’s t-test).

## *Msi1^+^* cells repopulate the intestinal epithelium independent of *Lgr5^high^* cells

To define the mechanism underlying +4 ISC-mediated epithelial regeneration, we also performed scRNA-seq on *Msi1^+^* cell progeny three and five days after irradiation. Three days after irradiation, considered as proliferative phase (Kim et al., 2017), ten distinct cell clusters were identified, including clusters of stem cells, EPs, EP-like cells, secretory precursors (SPs), ECs, goblet cells, EECs, tuft and Paneth cells (**Fig 5A and EV5A**). ISCs are subdivided into two clusters, IR3-C1 and IR3-C2. The first cluster is highly enriched for DDR genes (**Fig 5B**) and DNA helicases (**Figure EV5B**), and most cells are in S phase (**Fig 5C and EV5C**). Compared to IR3-C1, IR3-C2 cells are characterized by reduced levels of DDR and DNA helicase genes (**Fig 5B and EV5B**) and exhibit increased proliferative capacity, as evidenced by the enrichment for proliferating marker genes (**Figure EV5D**). Indeed, over 90% of IR3-C2 cells were in G2/M phase (**Fig 5C**). IR3-C3 cells localize in the EC branch before EP-like cells (**Fig 5D**), and most of them were in S and G2/M phases. Thus, IR3-C3 cells were identified as proliferating EPs. In comparison, IR3-C4 cells are dormant EP-like cells and are in G0/G1 phase (**Fig 5C**). Another important finding is that SPs (IR3-C6) start to emerge at this stage. The cells are defined by *Dll1* expression (van Es, Sato et al., 2012) (**Figure EV5E**), rapid proliferation (**Fig 5C**), and close relatedness to secretory differentiated cells in the pseudotime trajectory (**Fig 5D**) and on PCA analysis (**Fig EV5F**). Many proliferating goblet cells were identified three days after irradiation, compared to two days (**Fig 5A-C, and EV5C**). Interestingly, *Lgr5^high^* CBCs are not emerging at this stage (**Fig 5A and EV5G**), In agreement, immunohistochemical assay showed that the proportion of Lgr5^+^ cells is the lowest 3 days after irradiation (**Fig 5E and EV5H**). Thus, it appears that surviving ISCs directly give rise to proliferating EPs and proliferating SPs independent of *Lgr5^high^* cells.

**Figure 5.**
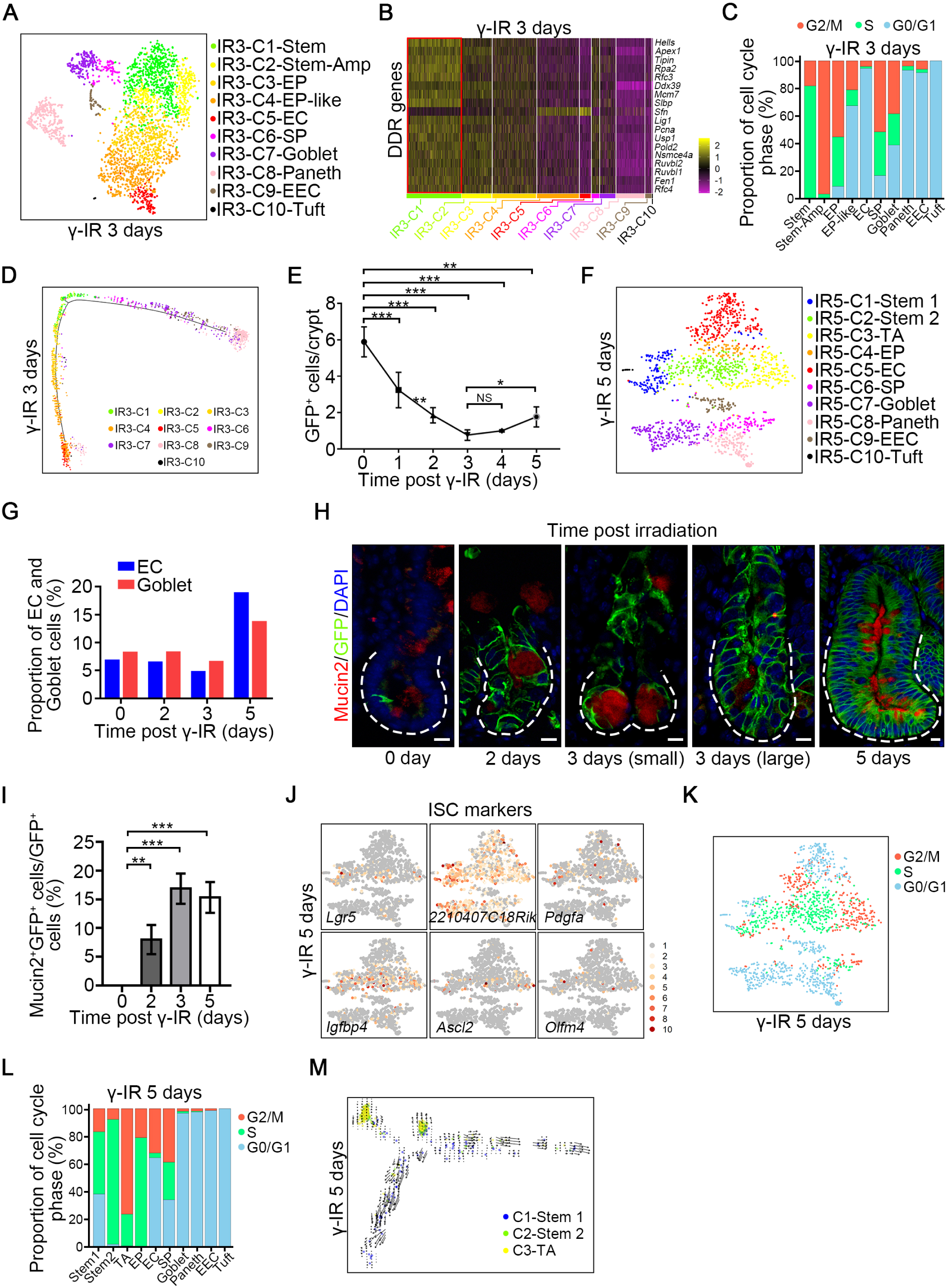
Surviving ISCs repopulate intestinal epithelium independent of CBCs. (A) A t-SNE plot revealed cellular heterogeneity with ten distinct clusters of *Msi1^+^* cell progeny from *Msi1^CreERT2^;R26^mTmG^* mice three days after γ-IR. The general identity of each cell cluster is defined on the right. (B) Heatmap of DDR genes in distinct clusters three days after γ-IR. (C) Proportion of cell cycle stages in each cluster three days after γ-IR. Cells in S phase are colored green, those in G2/M phase are red, and those in G0/G1 phase are blue. (D) Pseudotime ordering on *Msi1^+^* cell progeny three days after γ-IR. (E) Quantification of GFP^+^ cells in the intestinal crypts of *Lgr5^EGFP-IRES-CreERT2^* mice at the indicated time points after γ-IR. 180 intestinal crypts (60 crypts/mouse, n=3 mice) were quantified at each timepoint. Data represent the mean value ± SD. NS, not significant; \**P*<0.05; \*\**P*<0.01; \*\*\**P*<0.001 (Student’s t-test). Representative images were shown in Fig EV5G. (F) A t-SNE plot revealed cellular heterogeneity with ten distinct clusters of *Msi1^+^* cell progeny from *Msi1^CreERT2^;R26^mTmG^* mice five days after γ-IR. (G) The proportion of EC and goblet populations at the indicated time points after γ-IR. (H) Immunofluorescence for Mucin2 and GFP in *Msi1^CreERT2^;R26^mTmG^* normal intestinal crypts and regenerative foci at the indicated time points after γ-IR. “3 days (small)” indicates small regenerative foci; “3 days (large)” indicates large regenerative foci. Scale bar: 10 μm. (I) Quantification of the percentage of Mucin2^+^GFP^+^ cells versus GFP^+^ cells in Panel H (n=3 mice). \*\**P*<0.001; \*\*\**P*<0.001 (Student’s t-test). (J) Feature plots of expression distribution for ISC marker genes in t-SNE plots five days after γ-IR. Expression levels for each cell are color-coded. (K) Cell cycle metrics on *Msi1^+^* cell progeny five days after γ-IR. t-SNE plot of the assigned cell cycle stages on *Msi1^+^* cell progeny. Cells in S phase are colored green those in G2/M phase are red, and those in G0/G1 phase are blue. (L) Proportions of cell cycle stages in each cluster five days after γ-IR. Cells in S phase are colored green, those in G2/M phase are red, and those in G0/G1 phase are blue. (M) RNA velocity analysis of IR5-C1 to IR5-C3 across the pseudotime trajectory five days after γ-IR. In E and I, Data represent the mean value ± SD. NS, not significant; \**P*<0.05; \*\**P*<0.01; \*\*\**P*<0.001 (Student’s t-test).

Five days after irradiation, tissue enters the normalization phase (Kim et al., 2017), and dramatic changes were observed at this time in *Msi1^+^* progeny on scRNA-seq (**Fig 5F and EV5I**). Compared to three days after irradiation, the populations of EC and goblet cells expand dramatically (**Fig 5F and G**), while EP-like cells almost entirely disappear (**Fig 5F**). The increase in goblet cells was further confirmed by immunofluorescence (**Fig 5H and I**). Another striking finding was the emergence of a new type of stem cell (IR5-C1), which is very similar to the *Lgr5^high^* ISC population in physiology and is characterized by the enrichment of *Lgr5^+^, 2210407C18Ric^+^,* and *Pdgfa^+^* accompanied by the appearance of *Igfbp4^+^, Ascl2^+^* and *Olfm4^+^* (**Fig 5J**). Similar to homeostatic *Lgr5^high^* ISCs, a large number of IR5-C1 cells reside in G1 phase (**Fig 5K and L**). Furthermore, the RNA velocity (La Manno, Soldatov et al., 2018) on IR5-C1, C2 and C3 clusters revealed that IR5-C1 cells are likely derived from IR5-C2 and IR5-C3 cells (**Fig 5M and EV5J**). Together, our data indicate that new IR5-C1 cells are nascent *Lgr5^high^* ISCs. Overall, we posit that during epithelial regeneration, surviving ISCs directly give rise to proliferative precursors of differentiated lineages, and only later do they regenerate relatively slowly cycling *Lgr5^high^* ISCs. We conclude that the *Msi1^+^* cells repopulate the intestinal epithelium independent of *Lgr5^high^* cells.

### Msi1^+^ ISCs preferentially produce Paneth cells

Another striking finding that drew our attention was the dynamic change in Paneth cells during epithelial regeneration. Two days after irradiation, Paneth cells are the most abundant cell type, accounting for 39% (**Fig 6A and EV3D**), with this proportion decreasing to approximately 10% three to five days after irradiation (**Fig 6A**). Lineage-tracing analysis revealed a large number of *Msi1^+^* cell-derived Paneth cells (over 25%) residing in the regenerative unit two days after irradiation (**Fig 6B and C**). Three days after irradiation, there existed small and large regenerative units (**Fig 6B**). The proportion of *Msi1^+^* cell-derived Paneth cells is much higher in the small regenerative units than in the large ones (**Fig 6D**). On scRNA-seq, two days after irradiation, Paneth cells can be divided into three distinct clusters based on marker genes (**Fig 6E**). Compared with type 1 and type 2 Paneth cells, type 3 cells exhibit increased levels of *Gm14851* and *Defa22*, and reduced level of *Mptx2.* Expression levels of *AY761184* and *Defa3* appear to gradually increase from Paneth cell type 1 to type 3 (**Fig EV6A**). Type 1 Paneth cells, which were transcriptionally closest to goblet cells, gradually changed to type 2, and finally to type 3 (**Fig EV3C**). This finding suggests a gradual maturation process in the direction of Paneth cell type 1 to type 2 to type 3. We also noticed that Paneth cell markers, such as *Lyz1*, *Defa17* and *Gm15284* were expanded in the goblet cell population (**Fig 6E**), whereas they are usually specific for Paneth cells during homeostasis (**Fig EV2B**). At this stage, Paneth cells are preferentially generated relative to goblet cells. Paneth cells have been identified as a niche for ISCs under physiological conditions (Sato, van Es et al., 2011). Accordingly, the ISC ligand *Wnt3* was highly enriched in these Paneth cells (**Fig 6F**). Together, our findings indicate that *Msi1^+^* cells preferentially give rise to Paneth cells upon exposure to irradiation.

**Figure 6.**
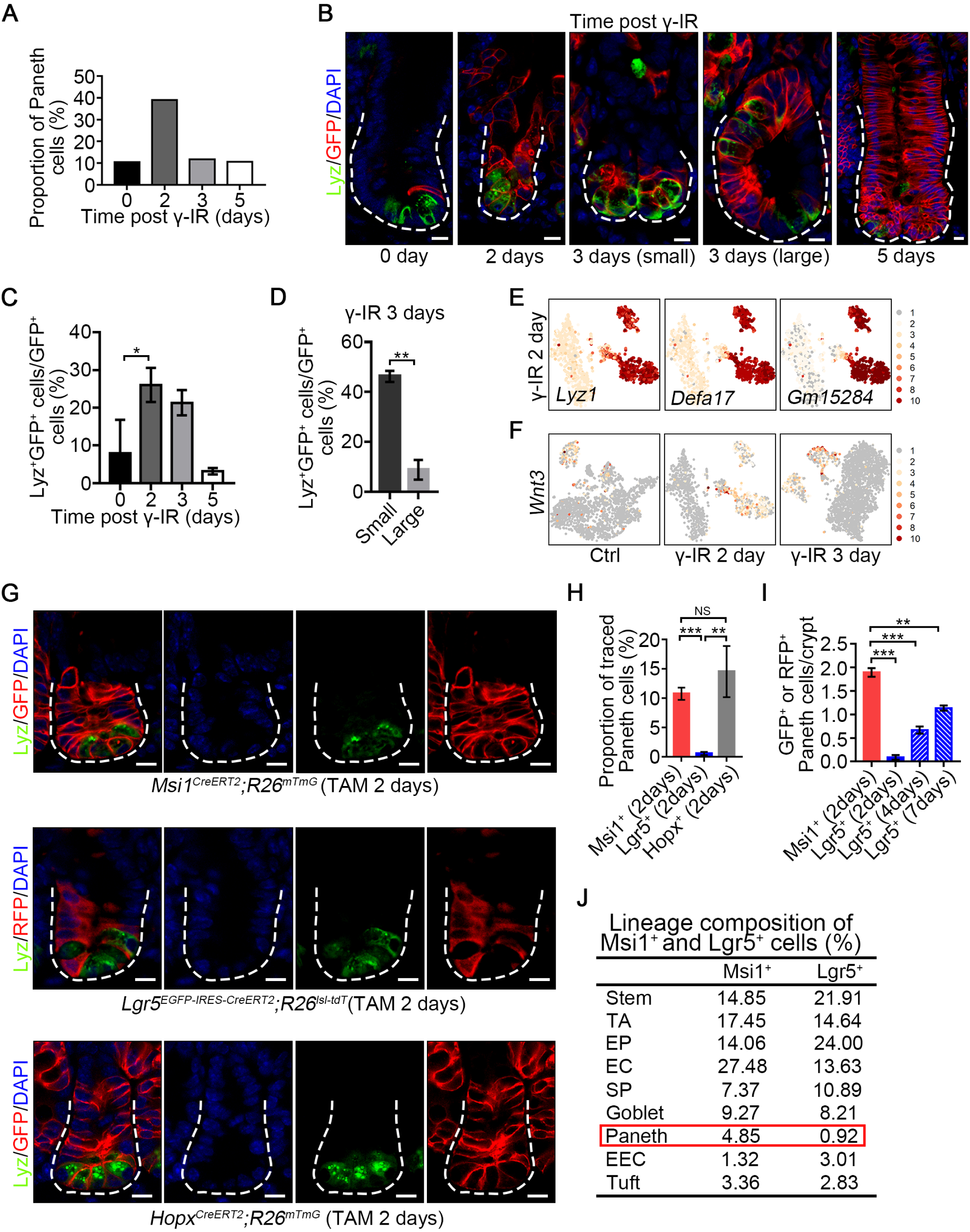
*Msi1^+^* ISCs preferentially generate Paneth cells two days after γ-IR. (A) The proportion of Paneth cells at indicated time points after γ-IR. (B) Immunofluorescence for lysozyme and GFP in normal intestinal crypts and regenerative foci from *Msi1^CreERT2^;R26^mTmG^* mice at indicated time points after γ-IR. “3 days (small)” indicates small regenerative foci; “3 days (large)” indicated large regenerative foci. Scale bar: 10 μm. (C) Quantification of Lyz^+^/GFP^+^ versus GFP^+^ cells (n=3 mice) at indicated time points in Panel B. \**P*<0.05 (Student’s t-test). (D) Quantification of Lyz^+^/GFP^+^ versus GFP^+^ cells in small (n = 133 crypts) and large (n = 66 crypts) regenerative foci three days after γ-IR. \*\**P*<0.01 (Student’s t-test). (E) Feature plots of expression distribution for Paneth cell marker genes two days after γ-IR. Expression levels for each cell are color-coded. (F) Feature plots of wnt3 distribution at indicated time points after γ-IR. Expression levels for each cell are color-coded. (G) Immunofluorescence for lysozyme and GFP/RFP in intestinal crypts from *Msi1^CreERT2^;R26^mTmG^*, *Lgr5^EGFP-CreERT2^;R26^lsl-tdT^* and *Hopx^CreERT2^;R26^mTmG^* mice two days after tamoxifen induction. Scale bar: 10 μm. (H) Quantification of Lyz^+^/GFP^+^ versus GFP^+^ (n=3 mice) or Lyz^+^/RFP^+^ versus RFP^+^ (n=3 mice) cells in Panel G. \*\**P*<0.01; \*\*\**P*<0.001; NS, not significant (Student’s t-test). (I) Quantification of the number of GFP^+^ or RFP^+^ Paneth cells in each crypt from *Msi1^CreERT2^;R26^mTmG^* and *Lgr5^EGFP-CreERT2^;R26^lsl-tdT^* mice at indicated timepoints after tamoxifen induction. n = 3 mice at each time point. Representative images are shown in Fig EV5B. \*\**P*<0.01; \*\*\**P*<0.001 (Student’s t-test). (J) scRNA sequencing revealed lineage composition of *Msi1^CreERT2^;R26^mTmG^* and *Lgr5^EGFP-CreERT2^;R26^lsl-tdT^* mice two days after tamoxifen induction. The t-SNE plots are shown in Fig EV5C and D. In C, D, H and I, data represent the mean value ± SD. \**P*<0.05; \*\**P*<0.01; \*\*\**P*<0.001 (Student’s t-test).

Considering the increase in *Msi1^+^* cell-derived Paneth cells two days after irradiation, we sought to examine whether *Msi1^+^* cells preferentially produce Paneth cells under normal physiological conditions. We quantified the number of Paneth cells after lineage tracing in *Lgr5^EGFP-IRES-CreERT2^;R26^lsl-tdT^* and *Msi1^CreERT2^;R26^mTmG^* mice two days after tamoxifen induction. Strikingly, we found that the proportion of *Msi1^+^* cell-derived Paneth cells is approximately 10.76% (60 crypts per mouse, n=3), while *Lgr5^+^* cell-derived Paneth cells are just 0.58% (50 crypts per mouse, n=3) (**Fig 6G and H**). We also observed that the proportion of *Lgr5^+^* cell-derived Paneth cells increased with the lineage tracing time (**Fig 6I and EV6B**), most likely due to the increase of +4 cells derived from *Lgr5^+^* cells with time. The finding of +4 cells preferentially generating Paneth cells were further confirmed by lineage tracing in *Hopx^CreERT2^;R26^mTmG^* mice two days after tamoxifen treatment (**Fig 6G and H**). These data suggest that *Msi1^+^* cells preferentially generate Paneth cells as compared to *Lgr5^+^* cells. To further confirm this idea, we performed scRNA-seq on labeled cells in intestinal crypt from *Lgr5^EGFP-IRES-CreERT2^;R26^lsl-tdT^* and *Msi1^CreERT2^;R26^mTmG^* mice two days after tamoxifen induction. Nine distinct cell clusters were identified in the progeny of *Msi1^+^* cells, and ten clusters in the progeny of *Lgr5^+^* cells. Consistently, we also found that the proportion of Paneth cells in *Msi1^+^* cell progeny is much higher than that of the *Lgr5^+^* progeny (**Fig 6J and EV6C-F**). In comparison, the proportions of goblet, tuft and EC cells were similar between them (**Fig 6J)**. Collectively, our findings strongly indicate that *Msi1^+^* cells preferentially produce Paneth cells during homeostasis relative to *Lgr5^+^* cells.

## Discussion

Our findings strongly indicate that the DNA damage-resistant subset of *Msi1^+^* ISCs, most likely *Lgr5^low/neg^* ISCs, are more rapidly cycling than *Lgr5^high^* CBCs (**Fig 7**), rather than quiescent, which substantially differs from the current intestinal stem cell theory. Classically, +4 cells have been identified as quiescent ISCs, while *Lgr5^high^* CBCs were thought to be rapidly cycling (Montgomery et al., 2011, Powell et al., 2012, Sangiorgi & Capecchi, 2008, Takeda et al., 2011, Yan et al., 2012). The notion of +4 ISCs dormancy was mainly supported by their co-localization with label-retaining cells in pulse-chase experiments. However, the +4 location of label-retaining cells (Potten et al., 1974, Potten et al., 2002) has been challenged by a number of subsequent studies. Three independent works demonstrated that the long-term label-retaining cells in intestinal crypts were Paneth cells and that short-term label-retaining cells were SPs undergoing commitment toward Paneth and EEC lineages (Buczacki et al., 2013, Li et al., 2016, Steinhauser et al., 2012). Likewise, *Bmi1*-expressing cells were recently identified as EEC lineage cells that possess ISC activity (Yan, Gevaert et al., 2017), although they were considered slow-cycling ISCs resistant to irradiation (Yan et al., 2012). The conclusion that label-retaining cells are terminally differentiated Paneth cells or SPs contrasts the notion that +4 cells are quiescent label-retaining ISCs. Our findings that *+* 4 cells cycle faster than *Lgr5^high^* cells is further supported by recent work, showing that *Lgr5^high^* CBCs are in an unlicensed G1 phase, while most cells in the +4 to +8 positions are in S phase (Carroll, Newton et al., 2018).

**Figure 7.**
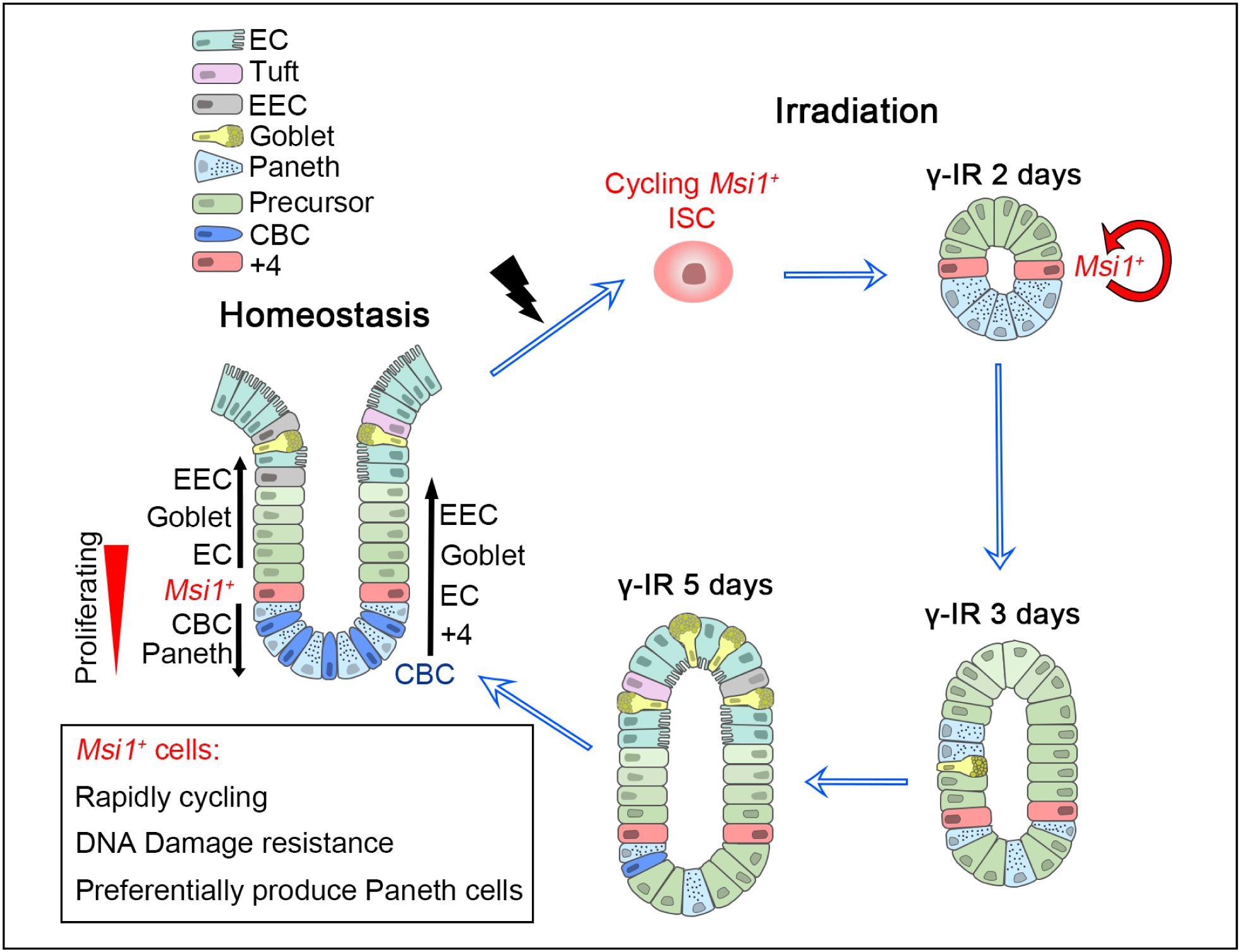
A model of *Msi1^+^* cells in maintaining and regenerating intestinal epithelium. A subset of *Msi1^+^* ISCs that exhibit DNA-damage resistance are cycling faster than *Lgr5^high^* CBCs, and regenerate the epithelium after DNA damage in a *Lgr5^high^* cell-independent manner. *Msi1^+^* cells preferentially produce Paneth cells during homeostasis and upon radiation repair.

Classic +4 ISC theory states that quiescent +4 cells become activated in response to irradiation. However, to the best of our knowledge, this idea lacks direct evidence. In contrast, it is well established that following radiation exposure, cells either transiently block cell cycle progression to allow time for repair, or exist cell cycle permanently (Shaltiel, Krenning et al., 2015). G1 arrest, S phase delay or G2 arrest can all take place following radiation-induced damage. Importantly, G1 arrest typically occurs at lower doses of irradiation, while S phase delay and G2 arrest are common at higher doses to allow for cells to repair DNA damage (Maity, McKenna et al., 1994). Accordingly, HR-mediated DNA repair, which enables an accurate repair using the sister chromatid as the template, can only occur in cycling cells during late S and G2 phases to repair DNA damage, making the cells survive from radiation exposure (Moynahan & Jasin, 2010). Another important factor in rendering S-phase cells resistant to DNA damage is that the signaling pathways regulating response to acute DNA damage also operate during normal S phase to maintain genome integrity in the presence of low levels of replication-associated damage (Ben-Yehoyada et al., 2007). Indeed, S-phase cells have been shown to be the least sensitive to irradiation (Pawlik & Keyomarsi, 2004). Consistently, we found that DNA damage repair genes are enriched in a subset of *Msi1^+^* cells during S and G2/M phases. Thus, we posit that the *Msi1^+^* ISCs during S and G2/M phase possess the capacity to resist DNA damage.

Although our data indicate that it is the rapidly-cycling *Msi1^+^* ISCs that survive irradiation exposure and repopulate damaged epithelium, we cannot formally rule out the previously proposed model that quiescent ISCs and/or precursors also contribute to epithelial regeneration (Ayyaz, Kumar et al., 2019, Chaves-Perez, Yilmaz et al., 2019, Yan et al., 2017). It is noteworthy that, while many secretory progenitor cells, marked by *Dll1-CreERT* (van Es et al., 2012), *Prox1-CreERT* (Yan et al., 2017), or H2B-label (Buczacki et al., 2013), have regenerative capacity, the contribution of these cells to epithelial regeneration are limited (Bankaitis, Ha et al., 2018). In comparison, the rapidly-cycling *Msi1^+^* ISCs might represent a primary source for regenerating intestinal epithelium. Furthermore, it is worth mentioning that the primary DNA damage repair pathway in quiescent stem cells in other tissues such as hematopoietic system—NHEJ—is error-prone, resulting in genome instability due to the accumulation of subtle mutations and chromosomal aberrations (Mohrin et al., 2010). If quiescent ISCs also use the same mechanism, many DNA mutations and chromosomal aberrations would exist in the surviving quiescent ISCs after radiation exposure. This would be detrimental to normal epithelial regeneration and would contribute to tumorigenesis. Therefore, we believe that cycling ISCs survive radiation exposure due to the high-fidelity HR-type repair.

Our data also demonstrate that the surviving *Msi1^+^* cells repopulate damaged intestinal epithelium in *Lgr5^high^* cell-independent manner at the early stage and give rise to nascent *Lgr5^high^* cells only at later time. This observation substantially differs from the prevailing idea that dormant surviving +4 cells function as reserve stem cells that upon activation generate rapidly cycling *Lgr5^high^* cells, that then go on to produce all differentiated lineages (Li & Clevers, 2010). Indeed, in our lineage studies, progeny of *Msi1^+^* cells can initially move both upwards and downwards the crypt relative to +4 position in normal physiology, suggesting that they could generate their progeny independent of *Lgr5^high^* CBCs during homeostasis. In agreement with our observation, classic cell migration tracing studies also demonstrated that all crypt cells ultimately derive from cells located at about the +4 position (Kaur & Potten, 1986, Potten, 1998, Qiu, Roberts et al., 1994). In other words, *Lgr5^high^* ISCs are not the only direct progeny of +4 ISCs. Thus, we posit that a subset of *Msi1^+^* cells are *bona fide* ISCs responsible for both normal homeostasis and epithelial regeneration, independent of *Lgr5^high^* ISCs (**Fig 7**).

## Materials and methods

### Mice

All mouse experiment procedures and protocols were evaluated and authorized by the Regulations of Beijing Laboratory Animal Management and were strictly in accordance with the guidelines of the Institutional Animal Care and Use Committee of China Agricultural University (approval number: SKLAB-2015-04-03). *Msi1^CreERT2^* mice were generated at the Model Animal Research Center of Nanjing University. *Lgr5^EGFP-IRES-CreERT2^*, *R26^mTmG^*, *R26^tdT^* and *R26R^LacZ^* mice were purchased from Jackson Laboratories (stock number: 008875, 007676, 009427). *Hopx^CreERT2^* mice were obtained from John Epstein’s laboratory at the University of Pennsylvania. *R26^lsl--DTA^* mice were obtained from Sen Wu’s laboratory at China Agricultural University.

### Lineage tracing

For lineage tracing, eight-week-old mice were injected with a single pulse of tamoxifen (4 mg/25 g body weight, Sigma). To label the *Msi1^+^* cells at homeostasis, *Msi1^CreERT2^;R26^mTmG^* mice were administered with tamoxifen for fifteen hours before sacrifice. For the injury study, *Msi1^CreERT2^;R26R^LacZ^* and *Lgr5^EGFP-IRES-CreERT2^;R26R^LacZ^* mice were treated with 12 Gy γ-irradiation fifteen hours after a single pulse of tamoxifen, and sacrificed at indicated time points. In order to examine the survival after high doses of irradiation or cytotoxic damage, *Msi1^CreERT2^;R26^mTmG^*, and *Lgr5^EGFP-IRES-CreERT2^;R26^lsl-tdT^* mice were injected intraperitoneally with two doses of 5-FU within two days or 12 Gy γ-irradiation once and analyzed with FACS after two days. For cell proliferation assay, *Msi1^CreERT2^;R26^mTmG^* and *Lgr5^EGFP-IRES-CreERT2^;R26^lsl-tdT^* mice were intraperitoneally injected with EdU (0.2 mg/25 g body weight) for 1.5 hours before sacrifice.

To test whether S-phase *Msi1^+^* cells survived from exposure of 12Gy γ-irradiation, *Msi1^CreERT2^;R26^mTmG^* were pretreated with tamoxifen, intraperitoneally injected with EdU (0.017 mg/25 g body weight) 13.5 hours after tamoxifen induction, and then exposed to 12 Gy γ-irradiation 1.5 hours after EdU injection. The intestinal samples were harvested four hours, one day, two days and three days after exposure to γ-irradiation.

### LacZ staining

Tissues were fixed in fixative solution (0.2% glutaraldehyde, 5 mM EGTA, 2 mM MgCl_2_ in PBS) for two hours on ice, rinsed for ten minutes with detergent rinsing solution (0.02% NP40, 0.01% sodium deoxycholate, 2 mM MgCl_2_ in PBS) for three times and immersed in X-gal staining solution (5 mM K_3_Fe(CN)_6_, 5 mM K_4_Fe(CN)_6_, 0.02% NP40, 0.01% sodium deoxycholate, 2 mM MgCl_2_ 1 mg/mL X-gal in PBS) overnight at 37°C. The stained tissues were fixed in 4% paraformaldehyde (PFA) and dehydrated for paraffin embedding.

### Dual-staining for EdU and BrdU

Five-micrometer tissue paraffin sections were dewaxed, hydrated, incubated in 1 M hydrochloric acid at 37°C for twenty minutes, washed with PBS for three times and antigen retrieval was performed in 10 mM citric acid. The sections were then stained according to the manufacturer’s instructions using the Click-iT EdU Alexa Flour 594 kit (Beyotime, C0078S). After staining, the sections were incubated with blocking solution (Beyotime, P0102) for one hour at room temperature and incubated with primary antibody against BrdU (Abcam, ab6326, 1:100) overnight at 37°C. The sections were washed for three times, and incubated with 488-conjugated secondary antibodies (Thermo Fisher, A11006, 1:400) for one hour at room temperature, stained with DAPI for eight minutes, and finally mounted with anti-fluorescence quenching sealing medium.

### Histology, Immunohistochemistry (IHC) and Immunofluorescence (IF) assays

For histological staining, paraffin-embedded and 5-μm sections were stained with hematoxylin and eosin (H&E). Periodic acid-Schiff (PAS) staining was performed using standard methods. For immunohistochemistry staining, the sections were deparaffinized with xylene followed by treatment with serial dilutions of ethanol. Antigen-retrieval was performed by heating slides to 95°C for 10 min in 0.01 M citrate buffer (pH 6) in a microwave oven. After cooling to room temperature, sections were incubated with blocking solution for 1 hour after administration of 3% H_2_O_2_ to eliminate endogenous peroxidase activity. Then, the sections were incubated with primary antibody overnight at 4°C. The sections were then immunostained by the ABC peroxidase method (Vector Laboratories) with diaminobenzidine as the enzyme substrate and hematoxylin as a counterstain. For immunofluorescence staining, paraffin sections were microwave pretreated in 0.01 M citrate buffer (pH 6.0), and incubated with primary antibodies, then incubated with secondary antibodies (invitrogen) and counterstained with DAPI in mounting media. The primary antibodies included Ki67 (thermo fisher, RM-9106-S1,1:500), cleaved caspase 3 (CST, 9664s, 1:1000), lysozyme C (Santa Cruz, sc-27958, 1:500), ChgA (Abcam, ab15160, 1:400), Mucin2 (Santa Cruz, sc-15334, 1:500), pH3 (Abcam, ab14955, 1:200), BrdU (Abcam, ab6326, 1:100), PCNA (Abcam, ab92552, 1:200), GFP (Abcam, ab13970, 1:800), GFP (Abcam, ab290, 1:800), RFP (ROCKLAND, 600-401-379,1:200).

### Cell cycle calculation

Ten-week-old mice were intraperitoneally injected with EdU (0.2 mg/25 g body weight) 1.5 hours after a pulse of BrdU (1 mg/25 g body weight) and sacrificed 0.5 hours later. The calculation was based on the assumption that EdU and BrdU could not be detected within thirty minutes after administration into mice. Cells still in S phase during the labeled time were EdU^+^BrdU^+^ (S_cells_) whereas cells that had exited S phase were BrdU^+^ (L_cells_). The average cell cycle time (T_c_) and S phase length (T_s_) of +4 cells and CBCs were calculated according to the formulas below. The number of proliferating cells was calculated based on the percentage of PCNA^+^ cells in each stem cell in Figure 4*C*. T_j_ is the time during which cells can labeled with BrdU but not EdU.

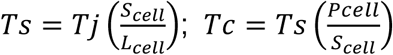

### In situ hybridization

The small intestine of 10-week-old mice was harvested and fixed in neutral buffered formalin (NBF) at room temperature (RT) for twenty four hours before paraffin embedding. The tissues were chopped into 5 µm sections and handled using Advanced Cell Diagnostics RNAscope 2.5 HD detection Reagents-RED kit (ACD) with mouse Msi1 probe (ACD, 469801). The detailed operation steps of *in situ* hybridization were followed according to the manufacturer’s instructions (322360-USM).

### Flow cytometry

The single-cell suspension of intestinal epithelium was collected as described previously (Sato et al., 2009). The fresh mouse intestine was cut open longitudinally and the villi were scraped off. The tissue was chopped into 5 mm pieces and incubated with 10 mM EDTA in PBS for thirty minutes at 4°C. The crypt fractions were collected by pipetting and filtered through a 70 µm cell strainer (BD biosciences). The gathered crypts were centrifuged at 1200 rpm for five minutes and digested with dispase (1 U/ml, Stem Cell Technologies). The single cell suspension was passed through a 40µm cell strainer (BD biosciences) and stained with Fixable Viability Dye (eBioscience, 65-0863-14) to remove dead cells. The flow cytometry analysis was performed on a BD FACS Arial 3.0. *Msi1^+^* cells were quantified by cells separated from *Msi1^CreERT2^; R26^mTmG^* mice 15 hours after tamoxifen induction. *Lgr5^high^* cells were sorted by flow cytometry from *Lgr5^EGFP-IRES-CreERT2^* mice.

### qRT-PCR analysis

All collected cells were sorted into TRIzol (Thermo Fisher, 10296010) immediately and total RNA was extracted using the RNeasy Plus Mini Kit (Qiagen, 74134). Real-time PCR was performed on a LightCycler 480 real-time PCR system (Roche) combined with the LightCycler 480 SYBR Green I master mix (Roche, 04887352001). The primers used for the gene expression assessment were as follows:

*Olfm4*-forward, 5’-CAGCCACTTTCCAATTTCACTG-3’; *Olfm4*-reverse, 5’-GCTGGACATACTCCTTCACCTTA-3’;

*Lgr5*-forward, 5’-CCTACTCGAAGACTTACCCAGT-3’; *Lgr5*-reverse, 5’-GCATTGGGGTGAATGATAGCA-3’;

*Axin2*-forward, 5’-TGACTCTCCTTCCAGATCCCA-3’; *Axin2*-reverse, 5’-TGCCCACACTAGGCTGACA-3’;

*Sox9*-forward, 5’-GCAGACCAGTACCCGCATCT-3’; *Sox9*-reverse, 5’-CGCTTGTCCGTTCTTCACC-3’;

*Ascl2*-forward, 5’-AAGCACACCTTGACTGGTACG-3’; *Ascl2*-reverse, 5’-AAGTGGACGTTTGCACCTTCA-3’;

### Single-cell mRNA sequencing

A single-cell suspension of intestinal epithelium was prepared as described above. The cells were stained with Fixable Viability Dye (eBioscience, 65-0863-14), CD45 (eBioscience, 17-0451-82), CD31 (eBioscience, 17-0311-82), TER119 (eBioscience, 17-5921-82), to remove dead and lin^-^ cells, and GFP^+^ cells were sorted into EP tubes in single-cell mode by FACS. The collected cells were held on ice before loaded for GemCode single cell platform (10X). Chromium Single Cell 3’ v2 libraries were sequenced with a Novaseq 6000 sequencer, with the following sequencing parameters: read 1, 150 cycles; i7 index, 8 cycles and read 2, 150 cycles.

### Primary computational analysis

Raw Illumina data were demultiplexed and processed using Cell Ranger (10X Genomics version Cell Ranger 2.0.1). The MM10 reference transcriptome provided by 10X genomics was used for mapping. Seurat version 2.3.4 was used for filtering and subsequent clustering (Butler, Hoffman et al., 2018). In order to remove partial cells and doublets, cells with less than 1000 genes or more than 7000 genes were removed. Additionally, cells with more than 10% of mitochondrial unique molecular identifiers (UMIs) were removed, as a high proportion of mitochondrial expression in cells is indicative of cell stress/damage during isolation. In order to reduce gene expression noise, genes that are expressed in 6 cells or less are removed. Gene-cell matrices were normalized and scaled in Seurat using default parameters for UMIs. Highly variable genes were found using a lower x threshold of 0.0125 and a y threshold of 0.5. Principal Component Analysis (PCA) was performed using the highly variable genes identified. T-distributed stochastic neighbor embedding (t-SNE) was performed using the PCA reduction. PCA reduction was also used to clusters with standard modularity function. Because of their low numbers, tuft cells in each time point were manually identified based on expression of canonical markers. A likelihood-ratio test for single cell gene expression was used to identify marker genes for each population (McDavid, Finak et al., 2013). Single-cell consensus clustering (SC3) analysis was used to validate the robustness of some clusters (Kiselev et al., 2017). Cell cycle analysis was carried out in Seurat using a list of cell cycle genes from the Regev laboratory (Kowalczyk, Tirosh et al., 2015).

### Pseudotime

Monocle version 2.10.1 was used on cells filtered from Seurat to infer differentiation trajectories (Qiu et al., 2017). An expression threshold of 0.1 was applied. The highly variable genes identified from Seurat were used as the ordering filter. DDRTree was used for dimension reduction. Initially, no root state was specified and the cells were ordered in an unsupervised manner. After the trajectory was obtained, a root state was specified based on where the stem cell populations are for subsequent systematic identification of pseudotime-dependent genes.

### Identification of pseudotime-dependent gene dynamics

We performed scEpath (Jin, MacLean et al., 2018) on Monocle-ordered cells to identify pseudotime-dependent gene expression changes as before (Guerrero-Juarez, Dedhia et al., 2019). Briefly, we compared the standard deviation of the observed gene expressions by randomly permuting the cell order (*nboot* = 100 permutations). Genes with a standard deviation greater than **0.5** and a Bonferroni-corrected *p-* value below a significance level ***α* = 0.01** were considered to be pseudotime-dependent. Pseudotime-dependent mouse transcription factors were annotated using the Animal Transcription Factor Database (AnimalTFDB 2.0).

## Acknowledgements

Z.Y. is funded by grants from the National Natural Science Foundation of China (81772984, 81572614); the Major Project for Cultivation Technology (2016ZX08008001, 2014ZX08008001); Basic Research Program (2015QC0104, 2015TC041, 2016SY001, 2016QC086); SKLB Open Grant (2018SKLAB6-12). C.F.G-J. is supported by the University of California Irvine Chancellor’s ADVANCE Postdoctoral Fellowship Program. ZL is supported by NIH T32-Training Program in Cancer Biology and Therapeutics. BA is supported by RO1AR44882.

## Author Contributions

ZY and FR designed research; XS, CL, CS, XB, MD, JX, ML, XW, QW and RZ performed research; XS, ZL, CFG-J, XL, QN, WC, SG, HZ, ZL, MP, CJL, BA, FR and ZY analyzed data; XS, CL and ZY wrote the manuscript.

## Conflict of interest

The authors have declared that no conflict of interest exists.

## Data availability

We are uploading the raw data of scRNA-sequencing, and will be available soon.

## Expanded View figure legends

**Figure EV1. Generation of *Msi1-CreERT2* knockin mouse.**

(A) Immunohistochemistry for Msi1 in mouse intestinal crypts. n = 3. Scale bar: 10 μm. (B) *In situ* hybridization for *Msi1* in mouse intestines with RNAscope methods (left). * indicates cells highly expressing *Msi1* mRNA (*Msi1^high^*). Scale bar: 10 μm. Quantification of *Msi1^high^* cells at indicated position (Right). n = 142 crypts. (C) Schematic representation of the *Msi1^CreERT2^* targeting vector. (D) Quantification of LacZ^+^ clone size in intestinal crypts of *Msi1^CreERT2^;R26R^LacZ^* mice at the indicated chase time. n=195 images analyzed, 270-663 crypts per chase timepoint were analyzed. (E) LacZ^+^ clone frequency in intestinal crypts of *Msi1^CreERT2^;R26R^LacZ^* mice at the indicated chase timepoint. n = 195 images analyzed, 270-663 crypts per chase timepoint were analyzed. (F) Flow cytometry analysis of intestinal epithelial cells from *Lgr5^EGFP-IRES-CreERT2^* mice, and in sorted *Msi1^+^* and *Hopx^+^* cells in *Msi1^CreERT2^;R26^mTmG^* and *Hopx^CreERT2^;R26^mTmG^* mice fifteen hours after tamoxifen induction. (G) qRT-PCR analysis for *Lgr5*, *Olfm4*, *Ascl2*, *Axin2* and *Sox9* in sorted *Lgr5^high^* cells from *Lgr5^EGFP-IRES-CreERT2^* mice and in sorted *Msi1^+^* and *Hopx^+^* cells in *Msi1^CreERT2^;R26^mTmG^* and *Hopx^CreERT2^;R26^mTmG^* mice fifteen hours after tamoxifen induction, as shown in Panel F. n = 3. \**P*<0.05; \*\**P*<0.01 (Student’s t-test). (H) Representative images of GFP^+^ ribbons in *Msi1^CreERT2^;R26^mTmG^* lineage-labeled small intestines four and six days after tamoxifen induction, or the mice were irradiated after fifteen-hour tamoxifen exposure, and harvested three and five days after γ-IR. Scale bar: 50 μm. (I) Quantification of regenerative foci in main Fig 1H.

In G and I, data represent the mean value ± SD. \**P*<0.05; \*\**P*<0.01; \*\*\**P*<0.001 (Student’s t-test).

**Figure EV2. Quality control metrics of scRNA-seq analysis from *Msi1^+^* cells.**

(A) Uniqe molecular identifiers (UMIs): 28753 UMIs per cell, 4316 genes per cells, The green line mark cutoffs for selecting cells. (B) Mean expression (log_2_(TMP+1)) of several marker genes for a particular cell type shown on t-SNE plots. (C) Feature plots of expression distribution for ISC marker genes, *Jun*, *Olfm4* and *Gkn3*. Expression levels for each cell are color-coded. (D) Expression levels of the ISC marker *Igfbp4*, the EC marker *Apoa1*, the goblet cell marker *Agr2*, the Paneth cell marker *Defa17*, the tuft cell marker *Lrmp* and the EEC marker *Neurod1* shown as a pseudotime feature plot. Expression levels for each cell are color-coded.

**Figure EV3. Rapidly cycling ISCs initiate intestinal epithelial regeneration.**

(A) Immunohistochemistry for Ki67 in the intestine at the indicated time points after γ-IR. Scale bar: 20 μm. (B) Quantification of Ki67^+^ cells per crypt in Panel A. NS, not significant; \*\*\**P*<0.001 (Student’s t-test). (C) Heatmap of differentially expressed genes in each cluster two days after γ-IR. (D) Mean expression (log_2_(TMP+1)) of several marker genes for a particular cell type shown on t-SNE plots, the selected marker genes were identical to Fig EV2B. (E) *Msi1^+^* cells are more radioresistant to DNA damage stress than *Lgr5^+^* cells. Mice were treated with γ-IR or two consecutive doses of 5-FU and then induced by tamoxifen fifteen hours before sacrifice. The quantification results were shown in Fig 3C. (F) Heatmap of cell survival genes in distinct clusters two days after γ-IR. (G) Violin plots of proliferating marker score in distinct clusters two days after γ-IR. (H) A t-SNE plot revealed cellular heterogeneity with ten distinct clusters of *Msi1^+^* cell progeny from *Msi1^CreERT2^;R26^mTmG^* mice one day after γ-IR. The mice were pretreated with tamoxifen fifteen hours before irradiation. General identity of each cell cluster is defined on the right. (I) Cell cycle metrics on *Msi1^+^* cell progeny one day after γ-IR. t-SNE plot of assigned cycling stages on *Msi1^+^* cell progeny. Cells in S phase are colored green, those in G2/M phase are red, and those in G0/G1 phase are blue. Data represent the mean value ± SD. NS, not significant; \*\*\**P*<0.001 (Student’s t-test).

**Figure EV4. *Msi1^+^* cells are more rapidly cycling compared to CBCs.**

(A) Immunofluoresence for EdU in intestinal crypts after a 90 min pulse of EdU at the indicated concentration. Scale bar, 10 μm. (B) Feature plots of expression distribution for DDR genes in *Msi1^+^* cells at homeostasis. Expression levels for each cell are color-coded. (C) Heatmap of genes functioning on cell survival and stress in distinct clusters. (D) Feature plots of expression distribution for the key genes functioning in HR-type DNA damage repair one days after irradiation. Expression levels for each cell are color-coded.

**Figure EV5. scRNA-sequencing analysis on *Msi1^+^* cell progeny three and five days after γ-IR.**

(A) Heatmap of differentially expressed genes in distinct clusters three days after γ-IR. (B) Heatmap of DNA helicase genes in distinct clusters at the indicated time points after γ-IR. (C) Cell cycle metrics of *Msi1^+^* cell progeny three days after γ-IR. t-SNE plot of the assigned cell cycle stages on *Msi1^+^* cell progeny. Cells in S phase are colored green, those in G2/M phase are red, and those in G0/G1 phase are blue. (D) Violin plots of proliferating marker gene score at indicated time point after γ-IR. (E) Feature plots of expression distribution for secretory progenitor marker *Dll1*. Expression levels for each cell are color-coded. (F) PCA analysis showing the association among distinct clusters three days after γ-IR. (G) Feature plots of expression distribution for CBC marker genes. Expression levels for each cell are color-coded. (H) Immunohistochemistry for GFP in the intestine of *Lgr5^EGFP-IRES-CreERT2^* mice at the indicated time points after γ-IR. Scale bar: 20 μm. (I) Heatmap of differentially expressed genes in distinct clusters five days after γ-IR. (J) RNA velocity analysis of IR5-C1 to IR5-C3 across the PCA plot five days after γ-IR.

**Figure EV6. *Msi1^+^* cells preferentially generate Paneth cells.**

(A) Feature plots of expression distribution for Paneth cell marker genes two days after γ-IR. Expression levels for each cell are color-coded. (B) Immunofluorescence for lysozyme and RFP in intestinal crypts from *Lgr5^EGFP-IRES-CreERT2^;R26^lsl-tdT^* mice four and seven days after tamoxifen induction. Scale bar: 10 μm. (C) A t-SNE plot reveals cellular heterogeneity, with nine distinct clusters of *Msi1^+^* cell progeny from *Msi1^CreERT2^;R26^mTmG^* mice two days after tamoxifen induction. (D) A t-SNE plot revealed cellular heterogeneity with ten distinct clusters of *Lgr5^+^* cell progeny from *Lgr5^EGFP-IRES-CreERT2^;R26^lsl-tdT^* mice two days after tamoxifen induction. (E and F) Mean expression (log_2_(TMP+1)) of several marker genes for particular cell types in *Msi1^+^*(E) and Lgr5^+^ (F) cell progeny shown on t-SNE plots. The selected marker genes are identical to those in Fig EV2B.

